# An atlas of transcriptionally defined cell populations in the rat ventral tegmental area

**DOI:** 10.1101/2021.06.02.446737

**Authors:** Robert A. Phillips, Jennifer J. Tuscher, Samantha L. Black, Lara Ianov, Jeremy J. Day

## Abstract

The ventral tegmental area (VTA) is a complex brain region that is essential for reward function but is also implicated in neuropsychiatric diseases including substance abuse. While decades of research on VTA function have focused on the role of dopaminergic neurons, recent evidence has identified critical roles for VTA GABAergic and glutamatergic neurons in reward processes as well. Interestingly, molecular characterization has revealed that subsets of these neurons express genes involved in the transport, synthesis, and vesicular packaging of multiple neurotransmitters, providing evidence for co-release neurons. However, these studies have largely relied on low-throughput methods, and the molecular architecture of the VTA has not been comprehensively examined. Here, we performed single nucleus RNA-sequencing (snRNA-seq) on 21,600 VTA cells from male and female Sprague-Dawley rats to generate a transcriptional atlas of the rat VTA. We identified 16 transcriptionally distinct cell types within the VTA, including 7 neuronal populations. Further subclustering revealed several VTA neuronal populations expressing markers for more than one neurotransmitter system, with one cluster exhibiting high expression levels of genes involved in the synthesis and transport of GABA, glutamate, and dopamine. Finally, snRNA-seq enabled the de novo identification of thousands of marker genes for each transcriptionally distinct population, revealing cluster-specific enrichment of gene sets implicated in neuropsychiatric and neurodevelopmental disorders, as well as specific phenotypes associated with alcohol and tobacco use. Together, these results highlight the heterogeneity of cellular populations in the VTA and identify novel markers and disease-linked genes enriched in distinct neuronal subtypes.

## INTRODUCTION

Dopaminergic signaling within the mammalian brain is essential for reward learning, motivation, and motor coordination. The brain circuits critical for dopamine neurotransmission are highly conserved across vertebrate evolution, and dysregulation of this circuitry has been linked to substance abuse^1,2^, neurodevelopmental disorders, and neuropsychiatric disease^3–6^. For decades, the ventral tegmental area (VTA) has been studied for its role in dopaminergic neurotransmission and reward processes. The VTA is a complex brain region that sends dense projections of dopamine (DA) producing neurons to other reward-related areas of the brain, such as the nucleus accumbens (NAc) and prefrontal cortex (PFC). DA neuron firing and DA release in terminal regions dynamically encodes rewarding stimuli as well as cues that predict rewards^7–11^, and is thought to provide a learning signal consistent with reward prediction error computational functions^10,12,13^. Consistent with this role, DA neurotransmission is critical for reward-related behavioral conditioning^14,15^, and phasic activation of VTA DA neurons is sufficient for reward learning^16^.

Despite a heavy focus on dopaminergic neurotransmission in the VTA, there is also substantial evidence for the importance of VTA GABAergic and glutamatergic neurotransmission in reward-related behaviors. For example, the activation of µ-opioid receptors on GABAergic interneurons in the VTA results in the depolarization of DA neurons^17,18^. Furthermore, in the absence of DA release, optical stimulation of VTA glutamate neurons is sufficient for positive reinforcement^19^. This evidence suggests a role for multiple neurotransmitter systems in VTA function, which is further supported by molecular characterization studies that have identified heterogeneous populations of dopaminergic, glutamatergic, and GABAergic neurons. These studies have also identified spatial heterogeneity in neuronal subtypes across the VTA. Specifically, DA neurons (neurons expressing genes involved in both the synthesis and transport of DA) are concentrated in the lateral area of the VTA, while glutamate neurons (expressing VGLUT2 mRNA) but not protein for tyrosine hydroxylase (TH, a common DA neuron marker), are concentrated in the medial portion of the VTA^20^. Interestingly, parallel lines of evidence have demonstrated the presence of “combinatorial” VTA neurons that co-express genes involved in the synthesis and transport of multiple neurotransmitters^21–24^, suggesting a new layer of complexity in cellular and synaptic function in the VTA^25,26^.

Previous studies investigating the molecular heterogeneity of cell types within the VTA have typically relied on low-throughput methods that do not provide comprehensive information on transcriptional diversity and heterogeneity within this brain region. Recent advances in single cell RNA-sequencing (scRNA-seq) technologies have allowed for the interrogation of whole transcriptomes from many cells within the VTA, circumventing previous technical limitations. These studies have provided key insights into transcriptomic diversity within VTA cell types^27^, DA neuron development^28^, as well as VTA cell type composition similarities between species^29^. However, these studies have either focused on a single cell type using a previously defined marker gene or combined other midbrain regions together for analysis. Furthermore, these studies have only used mice and have not included sex as a biological variable. Here, we performed single nucleus RNA-sequencing (snRNA-seq) on 21,600 VTA nuclei from both male and female adult Sprague-Dawley rats, a commonly used model organism for studies of reward function and substance abuse. Investigation of neuronal subpopulations confirmed the presence of combinatorial neurons and identified novel marker genes for newly characterized combinatorial neurons as well as classically-defined DA neurons. Moreover, we demonstrate cell type specific enrichment for gene sets implicated in multiple genome-wide association studies (GWAS) for neurodegenerative, neuropsychiatric, and neurodevelopmental diseases, as well as addiction-related phenotypes. This atlas, available as a searchable online resource (www.day-lab.org/resources), highlights novel avenues for regulation and identification of selected VTA cellular populations.

## RESULTS

### Identification of 16 Transcriptionally Distinct Cell Populations in the Rat VTA

To investigate cellular heterogeneity within the VTA, we used the 10x Chromium platform to perform snRNA-seq on 21,600 individual VTA nuclei from male and female adult rats (Fig. 1a). Following data integration, unsupervised dimensionality reduction techniques revealed 16 transcriptionally distinct cell populations (Fig. 1b-h and S1), which were equally represented in both sexes (Fig. S2). These cell populations included well-characterized glial cell types as well as VTA DA neurons that selectively express *Th* and *Slc18a2* mRNA, genes encoding TH and the vesicular monoamine transporter 2 (VMAT2) (Fig. 1b-d and Table S1). Th and *Slc18a2* were chosen as markers of DA neurons, as cells that selectively express these genes have the molecular machinery required to both synthesize and package DA into vesicles for release^21,30,31^.

**Figure 1.**
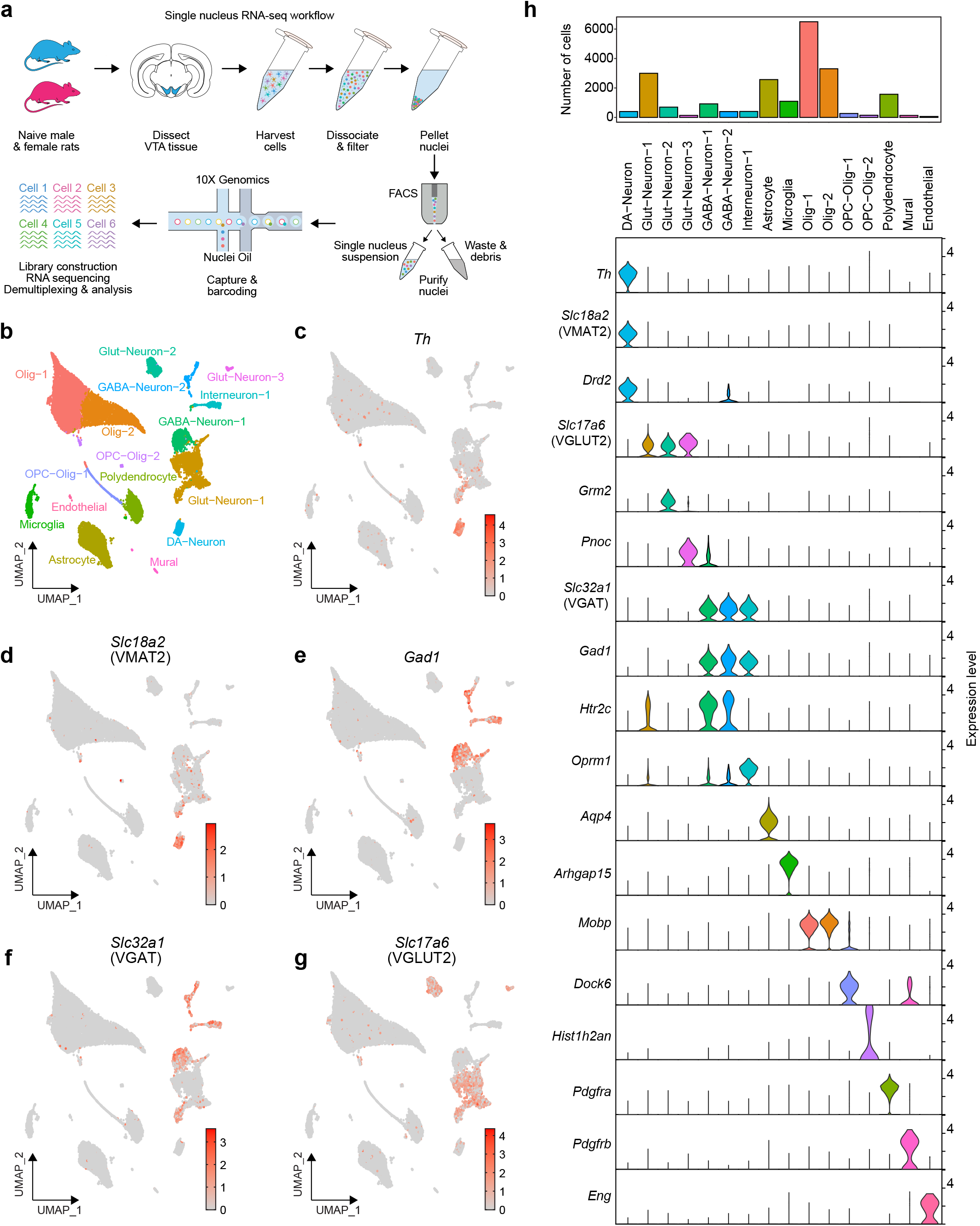
Transcriptional atlas of cell types in the rat ventral tegmental area. **a**, Single nuclei RNA-seq workflow. VTA tissue was harvested from experimentally naive male and female rats (n=6/sex) prior to nuclei isolation and sequencing on the 10X Genomics platform. **b**, UMAP identifying cell types within the rat VTA. **c-g**, Feature plots of expression values for major marker genes of dopaminergic, glutamatergic, and GABAergic neurons within cell types in the VTA. Scaled expression is represented in color scale. **h**, Bar graph indicating number of cells in each cluster (top) and violin plots showing major marker genes for all clusters identified with snRNA-Seq (bottom).

A wealth of literature has elucidated mechanisms by which subsets of VTA GABAergic neurons modulate the activity of DA neurons, and this comprehensive snRNA-seq approach also provided the ability to simultaneously study the unique transcriptional architecture of these populations of cells in a single dataset. Interestingly, unsupervised dimensionality reduction also identified three populations of GABAergic neurons marked by high levels of expression of *Gad1*, the gene encoding glutamate decarboxylase 1 (*Gad1*), and *Slc32a1*, the gene encoding the vesicular GABA transporter (VGAT) (Fig. 1e-f). The Interneuron-1 cluster is a GABAergic interneuron population that exhibits high expression of *Oprm1*, the gene encoding the µ-opioid receptor (Fig. 1h). Agonists at this receptor produce hyperpolarization of this interneuron population and subsequent disinhibition of dopaminergic neurons, which may contribute to the reinforcing nature of opioids^17,18^. The remaining GABAergic neuron populations (GABA-Neuron-1 and GABA-Neuron-2) express *Htr2c*, the gene encoding the serotonin 2C receptor, and represent a previously identified cell population within the VTA^32^ (Fig. 1h).

Similar to the GABAergic heterogeneity identified in the VTA, unsupervised clustering techniques also identified three populations of glutamatergic neurons, each of which exhibited high expression of *Slc17a6*, the gene encoding the vesicular glutamate transporter 2 (VGLUT2) (Fig. 1g-h). While all three populations expressed *Slc17a6*, only one of these populations (Glut-Neuron-2) expressed *Grm2*, a group II metabotropic glutamate receptor that acts as an autoreceptor at excitatory synapses^33–35^ (Fig. 1h). Another one of these glutamatergic neuron populations (Glut-Neuron-3) exhibited high expression of *Pnoc*, the gene encoding a preproprotein for nociceptin (Fig. 1h). This neuronal population likely represents a recently characterized cell type that negatively regulates motivation for reward^36^. The third population of glutamatergic neurons expressing *Slc17a6* (Glut-Neuron-1) does not express *Pnoc* or *Grm2*. However, a small subset of cells within this population exhibited expression of *Th, Gad1, Slc18a2*, and *Slc32a1* (Fig. 1c-f). The presence of cells exhibiting co-expression of *Slc17a6* and genes involved in other neurotransmitter systems suggests the presence of neuronal subpopulations capable of co-release or co-synthesis of multiple neurotransmitters.

### Subclustering of Neuronal Cells Identifies Populations of Combinatorial Neurons

Emerging evidence suggests that within the VTA, small subsets of neurons can synthesize and release more than one neurotransmitter^21–23,37^. To explore neuronal populations within the VTA more thoroughly, we first subclustered all neuronal cells within the dataset and identified 11 transcriptionally distinct neuronal populations (Fig. 2a). With increased resolution to detect smaller neuronal populations, this analysis revealed several clusters expressing markers corresponding to more than one neurotransmitter system (Fig. 2b). To provide direct evidence for the presence of combinatorial neurons, we examined the proportion of cells within a single cluster co-expressing unique genes involved in the synthesis or release of GABA, glutamate, or DA (Fig. 2c). In this analysis, a neuronal population that uses a single neurotransmitter is expected to contain a significant number of cells co-expressing genes critical to the synthesis, packaging, and release of that neurotransmitter. We detected five neuronal clusters which appear to represent selective GABAergic (clusters 1, 4, and 6), glutamatergic (cluster 3), and dopaminergic neurons (cluster 5; Fig. 2b-c). However, ∼20% of the cells within each “selective” cluster co-express at least a single gene involved in another neurotransmitter system, suggesting inherent promiscuity in the co-expression of genes important for neurotransmitter synthesis, transport, and release. In contrast to these classically defined “single neurotransmitter” populations, we also detected additional clusters in which markers for multiple neurotransmitter systems were more abundantly co-expressed (clusters 2, 8, 9, and 10), which may represent neurons capable of combinatorial neurotransmitter release. For example, cluster 2 represents a population of glutamatergic neurons that co-expresses genes involved in the synthesis of DA and transport of GABA. Likewise, cells in cluster 8 expressed markers of GABAergic neurons, but also co-expressed *Slc17a6* mRNA. While clusters 2 and 8 primarily co-express marker genes involved in the synthesis or transport of two neurotransmitters, few cells within these clusters also express marker genes that would allow for the synthesis and transport of all three neurotransmitter systems. Interestingly, cells in Cluster 9 expressed at least one gene involved in the synthesis and transport of glutamate, GABA, and DA. However, these cells are devoid of *Grm2* and *Gad1* mRNA, two genes that are highly expressed in classically defined glutamatergic and GABAergic cell populations. Previous research has identified populations of neurons that co-express *Th* and *Slc17a6*^37,38^, *Th* and *Gad1/2*^24^, as well as *Th* and *Slc32a1* mRNA^24^. To our knowledge, the simultaneous co-expression of marker genes for all three neurotransmitter systems has not been identified.

**Figure 2.**
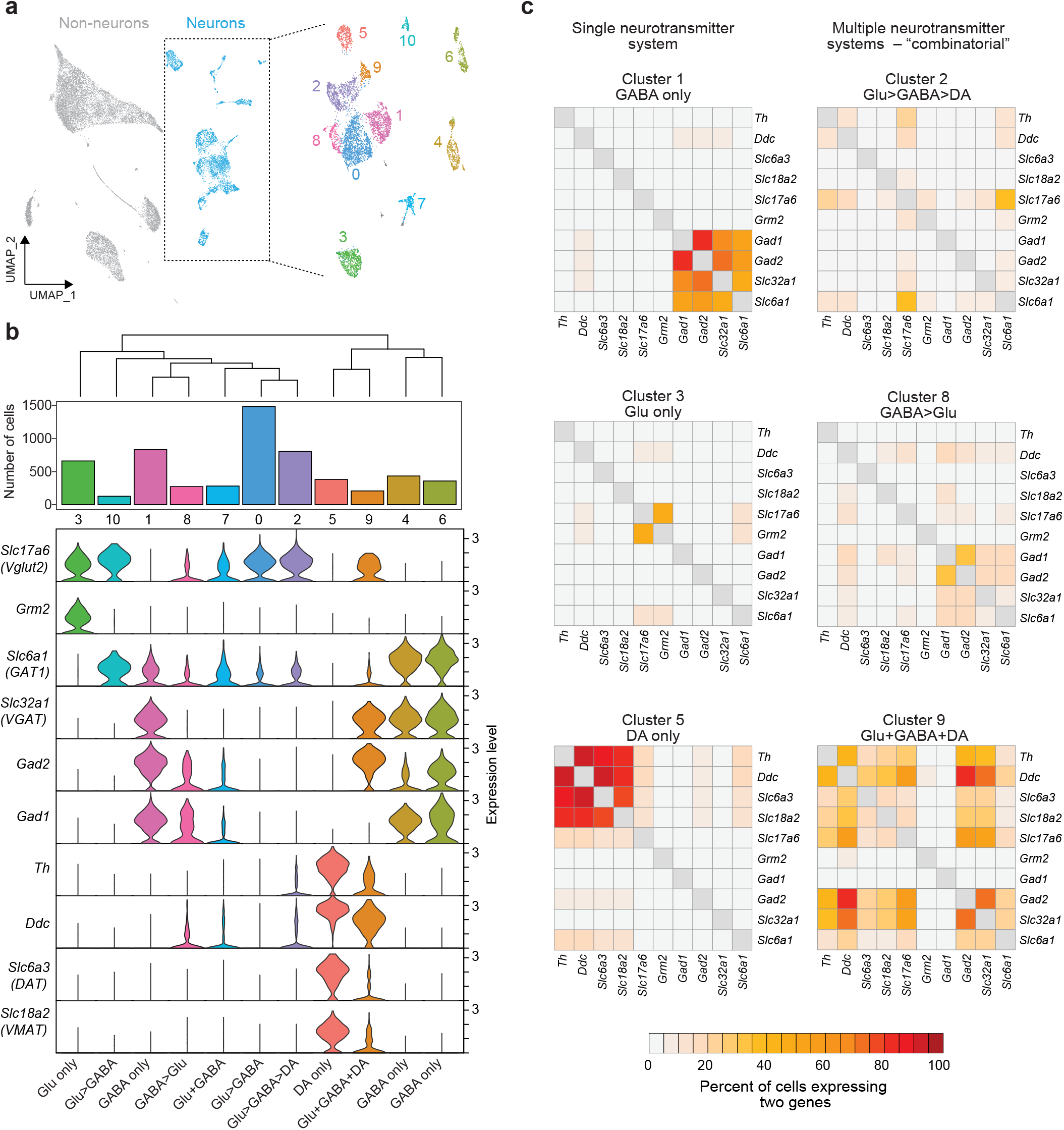
Transcriptional subclustering of VTA neurons reveals co-release and co-synthesis populations. **a**, Subclustering of neuronal populations identified 11 transcriptionally distinct cell types. **b**, Top, dendrogram and bar graph of VTA neuronal population numbers. Bottom, violin plots indicating expression values of genes involved in the synthesis and transport of DA, glutamate, and GABA. **c**, Heatmaps indicating the proportion of cells co-expressing genes involved in synthesis and transport of DA, glutamate and GABA.

### Neuronal Subcluster Analysis Identifies Novel Markers Genes

To identify novel markers of well-studied and potentially novel cell types in the VTA, we focused on neuron subcluster 5 (classically defined DA neurons) and neuron subcluster 9 (combinatorial neurons with high expression of genes involved in the synthesis and transport of GABA, glutamate, and DA). Using a Wilcoxon ranked-sum test^39^, we conducted differential expression testing to identify enrichment of genes in one neuronal subcluster versus all other neuronal clusters (Table S2). Classically defined DA neurons were enriched for genes involved in the synthesis and release of DA, such as *Th, Slc18a2* (VMAT2), and *Slc6a3* (DAT) (Fig. 3a and S3). While *Th* is widely used as a marker of DA neurons, only 36.8% of all *Th*-expressing neurons are found in cluster 5 (classically defined DA neurons). DA neurons were also enriched for *Dlk1*, a gene that encodes a transmembrane protein involved in the differentiation of DA neurons^40,41^, and Gch1, a gene encoding the enzyme GTP cyclohydrolase 1. Mutations in Gch1 have been implicated in genetic predisposition to dystonia and Parkinson’s diseases^42–45^, two diseases in which dysregulation of the DA neurotransmitter system has been extensively studied. Comparisons of DEGs for cluster 5 and cluster 9 found that classic markers of DA neurons, such as Th, *Slc18a2* (VMAT2), and *Slc6a3* (DAT) mRNA, were shared between the two clusters (Fig. 3b). While DEG testing allows for the identification of specific marker genes of neuronal subclusters, it does not provide any information for the inherent selectivity of gene expression patterns across all clusters. Thus, we employed the Gini coefficient, a statistic previously used to measure wealth inequality^46^, to characterize the selectivity of all detected genes. Here, a Gini coefficient of 0 would indicate that a gene is expressed in every neuronal subcluster equally, and a Gini coefficient of 1 would indicate that a gene is exclusively expressed in a single neuronal subcluster. The Gini coefficient for Gch1 is 0.76 (Fig. 3b), indicating that this gene exhibited a population-selective expression pattern. Furthermore, plotting the distribution of expression values for Gch1 across all neuronal subtypes revealed an enrichment within cluster 5 neurons (Fig. 3c), suggesting it is a selective marker of classically defined DA neurons. Similarly, differential expression testing identified *Slc26a7*, a gene encoding an anion transporter, as a marker for cluster 9 neurons (Fig. 3d). *Slc26a7* is required for chloride homeostasis in the stomach and kidney, but has been implicated in chloride transport in neurons in the olivocerebellar system^47^. Comparison of DEG lists identified that *Slc26a7* is a cluster 9 specific DEG (Fig. 3e). Furthermore, the high Gini coefficient for *Slc26a7* (0.79), combined with the high expression of this gene almost exclusively in cluster 9 neurons (Fig. 3f), indicated that this gene is a selective marker for this combinatorial neuron population.

**Figure 3.**
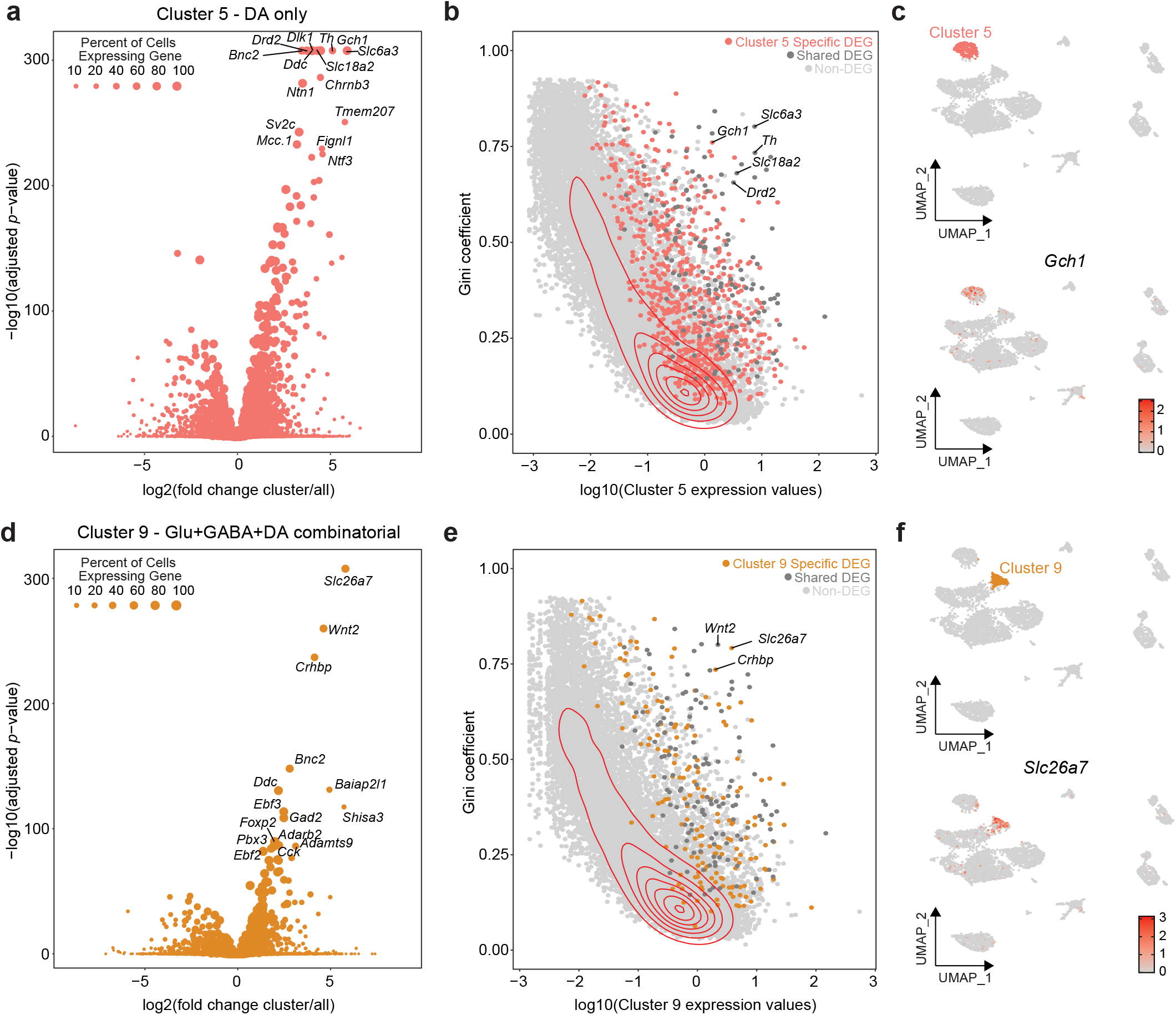
Identification of marker genes for classically defined dopamine and combinatorial neurons. **a**, Volcano plot depicting all DEGs for cluster 5 neurons. X-axis is log2 fold change between cluster 5 neurons and all other neurons. Top 15 genes by adjusted p-value are labeled. **b**, Scatter plot depicting the Gini coefficient and log transformed gene expression values for cluster 5 neurons. Colored points are cluster 5 specific DEGs, dark grey points are shared DEGs between cluster 5 and 9, and light grey points are non DEGs. **c**, UMAP depicting location of cluster 5 neurons and distribution of expression values for *Gch1*. **d**, Volcano plot depicting all DEGs for cluster 9 neurons. X-axis is log2 fold change between cluster 9 neurons and all other neurons. **e**, Scatter plot depicting the Gini coefficient and log transformed gene expression values for cluster 9 neurons. Colored points are cluster 9 specific DEGs, dark grey points are shared DEGs between cluster 5 and 9, and light grey points are non DEGs. **f**, UMAP depicting location of cluster 9 neurons and distribution of expression values for *Slc26a7*.

### Expression of DA Synthesis, Transport, and Degradation Genes

Although genes such as *Th, Slc6a3* (DAT), and *Slc18a2* (VMAT2) have all been used as markers of DA neurons, prior studies have not had the ability to systematically explore expression of all genes that regulate DA levels and localization in specific cell types. Therefore, we next focused on a set of genes known to play important roles in DA synthesis, transport, and degradation (Fig. 4). In the VTA, DA synthesis likely begins with tyrosine, a nonessential amino acid that crosses the blood-brain barrier. In the rate-limiting step in DA synthesis, tyrosine is converted to L-DOPA by tyrosine hydroxylase, using tetrahydrobiopterin (BH4) as a co-factor^48,49^. Notably, the GTP cyclohydrolase *Gch1*, identified in Fig. 3 as a selective marker of cluster 5 DA neurons, performs the rate-limiting step in BH4 synthesis by converting guanosine triphosphate (GTP) to dihydroneopterin triphosphate^50,51^ (Fig. 4b). This intermediary metabolite is further converted in a variety of de novo and salvage pathways to BH4, involving reductases such as sepiapterin reductase (*Spr*), aldo-keto reductase family 1, member B1 (*Akr1b1*), and dihydrofolate reductase (*Dhfr*)^50,52^. Surprisingly, in addition to *Gch1*, we observed significant enrichment of several other genes in the BH4 synthesis pathway in cluster 5 DA neurons, but not in cluster 9 “combinatorial” neurons that expressed other commonly used markers of DA synthesis and transport (e.g., *Th, Ddc, Slc6a3*, and *Slc18a2*). Taken together, these results suggest that combinatorial or “co-release” DA neurons have a paradoxically limited capacity for DA synthesis, despite the enrichment of *Th* and *Ddc* in this neuron cluster. Consistent with elevated DA synthesis capacity in cluster 5 DA neurons, we also observed enrichment of two genes involved in DA degradation (monoamine oxidase A, encoded by *Maoa*, and the previously identified DA neuron marker *Aldh1a1*, which codes for an aldehyde dehydrogenase) within this cluster (Fig. 4a-b).

**Figure 4.**
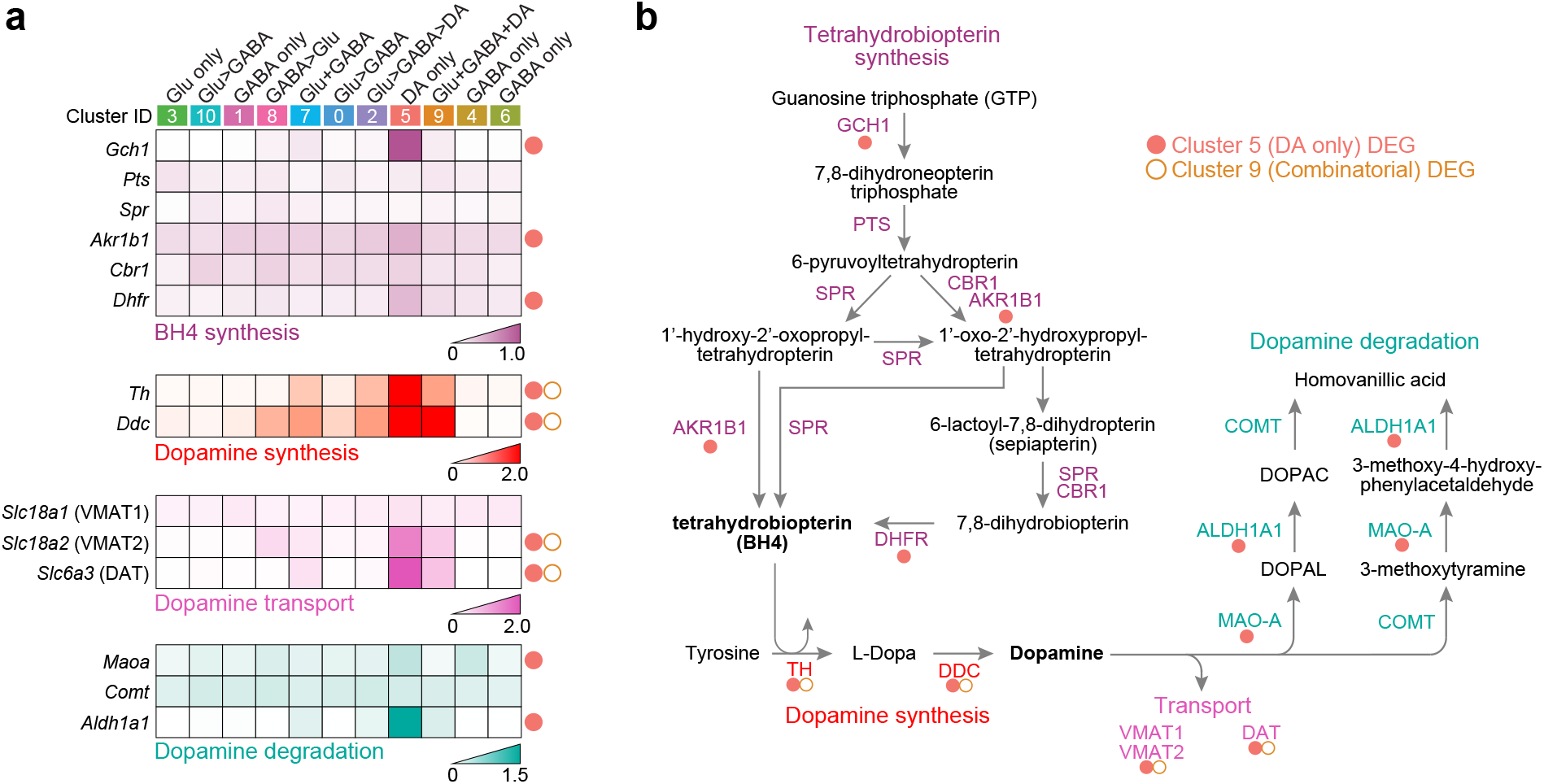
Expression of DA synthesis, transport, and degradation machinery in VTA neurons. **a**, Heatmaps showing mean expression values for genes involved in tetrahydrobiopterin (BH4) synthesis, DA synthesis, DA transport, and DA degradation. **b**, Pathway illustration for genes involved in BH4 and DA synthesis. Cluster 5 DA neurons express elevated levels of machinery for both DA and BH4 synthesis, as well as DA degradation.

### Opioid Neuropeptide Marker Analysis within Neuronal Subclusters of the VTA

Endogenous opioid neuropeptides and their receptors participate in a host physiological functions, including pain modulation, neuroprotection, euphoria, dysphoria, and maintenance of ionic homeostasis. However, our understanding of the cell-selective nature of opioid effects in the VTA remains poorly understood. To examine cell-type specific expression of opioid neuropeptide markers in the present dataset, we plotted the distribution of expression for several endogenous opioid peptide precursors (*Penk, Pomc, Pdyn, Pnoc, Tac1*), as well as the genes encoding their respective receptors (*Oprd1, Oprm1, Oprk1, Oprl1, Tacr1, Tacr3*; Fig. 5a). The pro-enkephalin gene *Penk*, a precursor for met- and leu-enkephalins implicated in analgesia, euphoria, and stress resilience, was highly enriched in the glutamatergic neuronal subcluster 3. In contrast, high affinity opioid receptors for met- and leu-enkephalins, *Oprd1* and *Oprm1*, exhibited limited expression in this glutamatergic subcluster, and were instead preferentially expressed in GABAergic cells in cluster 4, as well as other GABA subclusters and glutamate/DA combinatorial neurons to a lesser extent (Fig. 5a-b). This pattern of μ-opioid receptor expression is consistent with the excitatory actions of opioids in the VTA being mediated via GABAergic disinhibition of DA and glutamatergic neurons^17,19^. *Pomc*, the polypeptide precursor for several pro-opiomelanocortin derivatives including endorphins, melanotropins, and adrenocorticotropins, was present in all subclusters but at relatively low levels of expression. The nociceptin precursor *Pnoc* was also sparsely expressed in most clusters, with its highest enrichment in clusters 1 and 10 (classically defined GABA and Glu>GABA combinatorial neurons, respectively (Fig. 5a-b)). Similarly, the nociceptin receptor *Oprl1* was expressed in all neuronal subclusters, but to a limited extent. Tachykinin precursor gene *Tac1*, which is required for the synthesis of Substance P and neurokinins, was preferentially expressed in combinatorial subclusters for cluster 0 (Glu > GABA) and cluster 2 (Glu > GABA > DA), but was also present in clusters 4 and 6 (GABA only). Substance P receptor *Tacr1* was present in all clusters, but expressed at modest levels compared to neurokinin B receptor *Tacr3*, which was robustly expressed in the DA neuron cluster 5, as well as cluster 10 (Glu > GABA combinatorial neurons), and GABAergic clusters 4 and 6.

**Figure 5.**
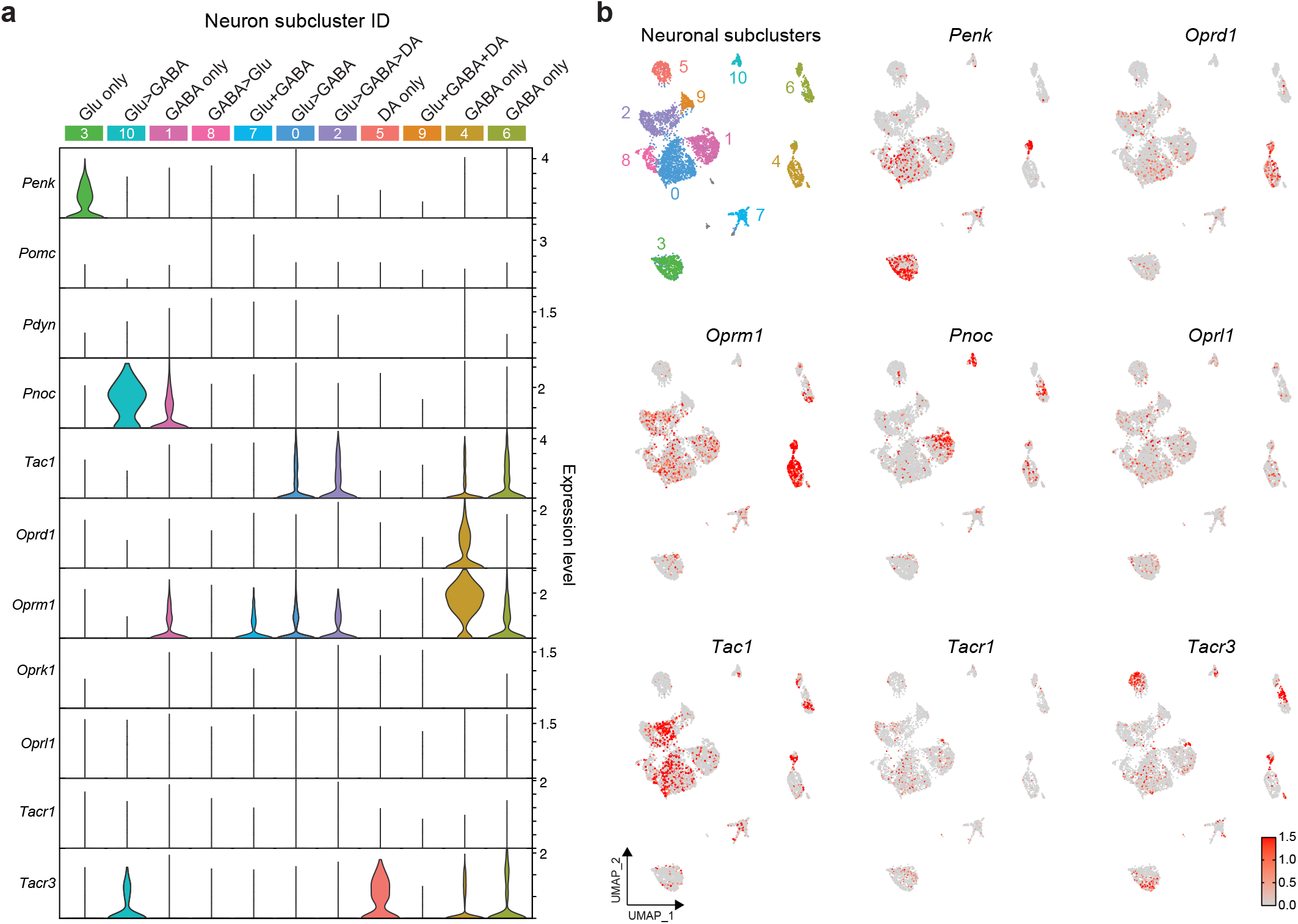
Endogenous opioid neuropeptides and their receptors are expressed in discrete neuronal sub-populations in the VTA. **a**, Violin plots indicating expression distribution of endogenous opioid precursor genes and their cognate receptors. **b**, Feature plots highlighting enrichment and distribution of selected opioid neuropeptide precursors and receptors across discrete neuronal subclusters.

### Cell Type Specific Gene-Set Enrichment for Brain Disorders and Addiction-Related Phenotypes

GWAS have identified risk genes that confer susceptibility to brain disorders and related phenotypes. Single cell sequencing enables further contextualization of these variants by identifying selected cell types that harbor enriched expression of risk genes. We noticed that expression of several genes implicated by GWAS to contribute to schizophrenia risk were unequally distributed across neuronal and non-neuronal cell types. For example, *Rims1*, a schizophrenia risk gene that encodes a protein responsible for the control of synaptic vesicle endocytosis, demonstrated a panneuronal enrichment (Fig. 6a). Therefore, we next sought to identify cell type specific enrichment for brain disorders and diseases with graphical analysis of risk gene enrichment in VTA cell types, as well as MAGMA^53^ (Multi-marker Analysis of GenoMic Annotation) gene set analysis. Specifically, we identified risk genes and mined summary statistics from GWAS and associated meta-analyses for Alzheimer’s disease^54^, Parkinson’s disease^55^, bipolar disorder^56^, schizophrenia^57^, attention deficit hyperactivity disorder (ADHD)^58^, autism spectrum disorders (ASDs)^59^, and specific traits associated with alcohol and tobacco use^60^.

**Figure 6.**
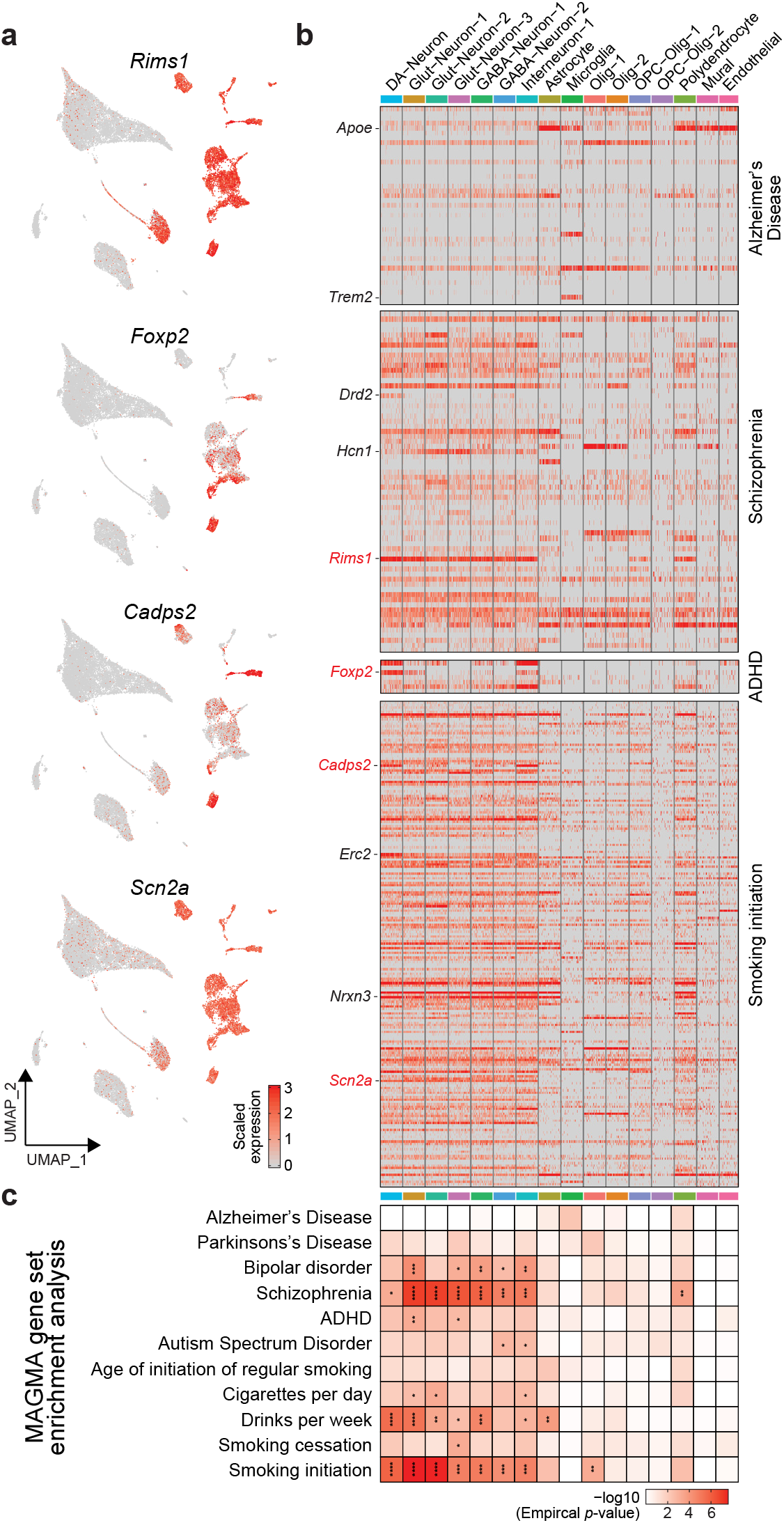
snRNA-seq from VTA reveals cell type specific enrichment for genes implicated in human neuropsychiatric and brain disorders by genome-wide association studies. **a**, UMAP expression plots for rat orthologues of selected human GWAS hits. **b**, Expression profiles of all rat homologs for human GWAS sets from Alzheimer’s Disease, Schizophrenia, ADHD, and smoking initiation. Scaled expression data was downsampled to 75 cells per cell type (columns), and rows represent individual genes. **c**, Heatmap of MAGMA analysis results depicting cell type specific gene set enrichment for neurodegenerative, neuropsychiatric, neurodevelopmental, and substance use phenotypes. Empirical p values are represented in color scale, and significant FDR-corrected observations are marked by asterisks (*adjusted p < 0.05, **p < 0.01, ***p < 0.001, ****p < 0.0001).

MAGMA gene set analysis identified a pan-neuronal association with SNPs identified in schizophrenia GWAS. Plotting the distribution of expression values for all schizophrenia genes identified a pan-neuronal enrichment (Fig. 6b). This enrichment of schizophrenia risk genes is expected given that the dysregulation of dopaminergic, glutamatergic, and GABAergic neurotransmission is heavily implicated in the development and progression of schizophrenia^61–63^. This analysis also identified a pan-neuronal enrichment for “smoking initiation”, a binary phenotype in which study participants were surveyed for whether they had ever been a regular smoker (Fig. 6b,c). Specifically, *Scn2a*, a gene which encodes a voltage-gated sodium channel was highly expressed in all VTA neurons. However, some genes such as *Cadps2*, a gene encoding a calcium binding protein responsible for the regulation of synaptic vesicle exocytosis, was most highly expressed in Glut-Neuron-1, DA neuron, and interneuron populations. The pan-neuronal enrichment of gene sets with SNPs identified in schizophrenia and smoking GWAS are interesting as upwards of 60% of schizophrenia patients are also smokers (Fig. 6c and Table S3). These findings suggest a shared genetic susceptibility within neuronal cell types of the VTA.

Consistent with the reported dysregulation of glutamatergic neurotransmission and signaling in ADHD^64–66^, we also found that ADHD risk genes were enriched in the Glut-Neuron-1 and Glut-Neuron-3 populations (Fig. 6c). Although dopaminergic signaling is also implicated in ADHD^67,68^ and the DA-neuron gene set was significantly enriched before multiple hypothesis testing correction, many of the combinatorial neurons that express genes involved in the synthesis and transport of DA originate from the Glut-Neuron-1 parent cluster. Taken together, these results demonstrate that specific populations of neurons within the VTA display an inherent genetic susceptibility to brain disorders and diseases that is most likely linked to their involvement in the synthesis and transport of specific neurotransmitters. These results also highlight the utility in using single nucleus transcriptomic technology to investigate the cell type specific expression of disease linked genes.

## DISCUSSION

The mammalian brain consists of complex, heterogenous regions that have become evolutionarily specialized for specific aspects of emotion, behavior, and homeostasis. This complexity is further compounded by the diversity of cell types within each region. Although the VTA has long been recognized as a site rich in cellular diversity, comprehensive molecular characterization of this region in the rat brain remains incomplete. Here, we present the first transcriptomic profiling atlas of cell types within the VTA of male and female Sprague-Dawley rats at single-cell resolution. Using unbiased dimensionality reduction approaches to analyze sequencing data from over 21,000 nuclei, we identified 16 transcriptionally distinct cell populations. Further subclustering of all major neuronal populations in this dataset revealed substantial cellular diversity, including rare combinatorial neurotransmitter populations with the potential for simultaneous release of GABA, glutamate, and DA. Neuronal subclustering also uncovered enrichment in gene markers for several opioid neuropeptide classes across 11 discrete neuron populations. Leveraging between-cluster statistical comparisons and measures of transcript selectivity, we next defined novel molecular markers for neuron subclasses, improving upon canonical DA neuron gene markers that are present in multiple populations. Finally, we used SNP-level gene set analysis to examine cell-type specific enrichment of GWAS risk-associated genes within the VTA, highlighting the mechanisms by which distinct cell populations may contribute to polygenic traits across several clinical disorders.

Although it is well established that the VTA is comprised of heterogenous cell types, historically, the majority of studies focused on this topic have relied on more targeted techniques, such as immunohistochemistry, in-situ hybridization, and single-cell RT-qPCR^21,22,37,69–71^. More recently, single-cell sequencing approaches have also been employed to investigate molecular heterogeneity within the VTA^27–30,72^. The present dataset expands on these studies in several key ways. For example, previous single-cell sequencing studies were conducted exclusively in the mouse brain^27–30,72^, and have relied primarily on sequencing a subset of FACS-isolated midbrain dopaminergic populations, rather than sampling all VTA cell types^28–30,72^. Here, we used an unbiased approach to conduct transcriptomic profiling of all cell types in the VTA, identifying 16 transcriptionally distinct cell populations and multiple neuronal subclasses not identified by earlier studies. Notably, this is also the first high resolution sequencing dataset to focus exclusively on VTA subregions, as other studies in mouse used tissues pooled from substantia nigra and VTA^27^, or a subset of fluorescently tagged FACS-isolated cells from general midbrain regions^28–30,72^. Finally, with over 21,000 nuclei from both male and female animals, our dataset also constitutes the largest and most comprehensive single-cell transcriptomic analysis focused exclusively on the cellular composition of the VTA.

The snRNA-seq approach utilized here afforded a previously unachieved level of molecular insight into the VTA, and underscores the transcriptional complexity within this brain region. Consistent with previous studies, we identified several neuronal populations with shared canonical GABAergic markers including *Gad1, Gad2*, and *Slc32a1* (VGAT)^21,32,73^. Unsupervised clustering allowed for further classification of GABAergic cells into GABA-Neuron-1, GABA-Neuron-2, and Interneuron-1 populations, each distinguishable by subclass-specific gene markers. Differential expression analysis revealed GABA-Neuron-1 and GABA-Neuron-2 share enrichment of the serotonergic receptor *Htr2c*, an observation consistent with previous research suggesting 5-HT2C receptors on GABAergic neurons contribute to inhibitory tone in the VTA^32^. In contrast, Interneuron-1 clusters contain limited *Htr2c* expression, and instead preferentially express the μ-opioid receptor *Oprm1*. Analysis of glutamatergic clusters in the VTA revealed similar transcriptional diversity. While all three glutamatergic populations were enriched with classical marker *Slc17a6* (VGLUT2), differential expression analysis identified subclass-defining markers unique to individual clusters. For example, the Glut-Neuron-2 cluster preferentially expressed metabotropic glutamate autoreceptor *Grm2*, whereas Glut-Neuron-3 cells exhibited enrichment for the nociceptin precursor *Pnoc*, a neuropeptide implicated in reward processing^36^. Interestingly, a subset of cells within the Glut-Neuron-1 population also expressed Th, *Gad1, Slc18a2*, and *Slc32a1* in addition to *Slc17a6*, suggesting a small subpopulation of these cells may be capable of co-release or co-synthesis of multiple neurotransmitters. Collectively, these observations support previous reports of high cellular diversity within GABA and glutamatergic populations of the VTA, and suggests discrete subclasses of these neurons may be modulated by distinct neurotransmitter and neuropeptide systems.

The presence of combinatorial neuronal populations with the capacity to regulate multiple neurotransmitter systems has been well-documented in the VTA^21–23,38,70,71,74^. Here we performed DEG analyses on 11 discrete neuronal subclusters to dissect the transcriptional complexity of co-release neurons. In addition to single neurotransmitter populations enriched with classical markers for GABA, glutamate, or DA regulation, we also detected several clusters harboring markers for the synthesis or transport of at least two different neurotransmitters. For example, subcluster 2 expressed classical glutamatergic markers, as well as *Th* and *Ddc*, genes involved in DA synthesis. Similarly, subcluster 8 contained canonical GABAergic markers in conjunction with dopaminergic marker *Ddc*. These observations are consistent with previous research identifying VTA combinatorial populations that co-express markers for both DA and glutamate^37,38^, or DA and GABA regulation^24^. Interestingly, although subcluster 9 was devoid of canonical glutamate and GABA markers *Grm2* and *Gad1*, this population co-expressed several markers indicative of the capacity for synthesis and transport of glutamate, GABA, and DA neurotransmitter systems. To our knowledge, this is the first report using an unbiased sequencing approach to identify neurons expressing the molecular machinery required to regulate all three neurotransmitter systems in a discrete population of cells.

Single cell sequencing enables robust differential expression testing to identify cell type specific genes that may be used for cell identification and viral targeting. Furthermore, differential expression testing can be used to identify new genes that can be used to target classically defined cell types, such as VTA DA neurons. Of the 21,600 nuclei captured, 399 exhibited gene markers for classically defined DA neurons. This proportion of DA neurons is in agreement with a previous scRNA-seq study that captured 991 DA neurons from a combined substantia nigra and VTA dataset of 44,091 cells^27^. Here, we focused on identifying marker genes for classically defined DA and combinatorial neurons. Identification of novel marker genes is important because genes widely used for the targeting of single neurotransmitter cell types, such as *Th* and *Slc17a6*, are also expressed in combinatorial neurons. Differential expression analysis identified *Gch1* as a novel, selective marker of classically defined DA neurons. *Gch1* encodes GTP cyclohydrolase 1, an enzyme that governs the rate limiting step in the synthesis of tetrahydrobiopterin, which acts as a co-factor for TH and is an essential molecule for the production for DA^44,75,76^. Thus, it is possible that cells lacking *Gch1* would not have the requisite machinery for de novo DA synthesis. Similarly, differential expression analysis also identified *Slc26a7*, a gene encoding an anion transporter, as a selective marker for combinatorial neurons with the capacity for GABA, glutamate, and DA synthesis. Together, these findings highlight the utility of leveraging snRNA-seq for molecular characterization and identification of novel, highly selective targets, as these genes have not previously been studied in the VTA and would not have been identified with previous methodologies.

In addition to classical DA, GABA, and glutamate neurotransmitter systems, the VTA is also a key site for neuropeptide actions. Opioid receptors have long been a focal point of neuropeptide research, and this focus has further intensified in response to the current opioid crisis. Here, we found that µ-opioid receptors were most abundant in GABAergic populations (Figs. 1&5). This is consistent with studies demonstrating that opioid-induced increases in DA and glutamatergic activity in the VTA are mediated predominantly by the disinhibition of GABAergic neurons^17,77^. In addition to robust expression in GABA populations, *Oprm1* is also moderately expressed in classically defined glutamate (Fig. 5, subcluster 3) and Glu/DA combinatorial neuron populations (Fig. 5, subcluster 2). This observation is in line with recent literature suggesting µ-opioid receptors may also regulate VTA neurotransmission by directly engaging glutamatergic neurons^19^. Further, the presence of these receptors in Glu/DA combinatorial neurons presents yet another means of fine tuning VTA neural activity and behavioral responses to opioids. Notably, expression of several opioid receptors (including *Oprd1, Oprm1*, and *Oprk1*) was low or absent in classically defined DA neuron populations, suggesting direct opioid modulation of DA neurons may be limited. Similarly, the pre-pronociceptin precursor encoded by *Pnoc* and its receptor *Oprl1* were also minimally expressed in DA neurons. Instead, neuronal subclustering revealed that while *Pnoc* is present in glutamatergic neurons, it is most enriched in GABAergic subclusters (Fig. 5, clusters 1, 4 & 6), as well as Glu/GABA combinatorial neurons (Fig. 5, cluster 10). This aligns with recent findings suggesting the VTA may contain two *Pnoc*^+^ subpopulations – one glutamatergic population anatomically centered in the ventromedial interpeduncular nucleus, and one GABAergic population in the lateral parabrachial pigmented nucleus^36^. Tachykinin precursors, including Substance P, neurokinins, and their respective receptors were also detected in VTA neuronal subclusters. Substance P, encoded by the pre-protachykinin gene *Tac1* and its receptor *Tacr1* mediate inflammatory response, vasodilation, and the perception of pain^78,79^. More recently, these genes have also been implicated in governing morphine-induced CPP and self-administration behaviors^80,81^, suggesting a role for tachykinins in opioid reinforcement. However, the exact mechanism through which these actions are regulated in the VTA remain unclear. For example, Substance P reportedly has excitatory effects on both DA and GABAergic neurons in the VTA^82^, but whether DA neuron activity is modulated directly or indirectly is not well established^83,84^. Neuronal subclustering in the present dataset suggests these effects may be predominately mediated through GABAergic or glutamate/DA combinatorial populations, rather than via direct modulation of DA neurons, where *Tac1* and its receptor *Tacr1* are largely absent. Overall, these findings confirm and extend previous research highlighting the complexity of neuropeptide modulation in multiple heterogenous neuron populations within the VTA.

While GWAS have begun to identify the genetic etiology of several brain disorders and diseases, these studies do not provide any evidence as to the specific cell type that could be susceptible to genetic disruption. To examine which cell types exhibit preferential enrichment of GWAS risk-associated genes, we conducted gene set analysis with MAGMA^53^, a multiple regression model that allows for assessment of the contribution of multiple gene markers to a particular clinical phenotype. This analysis identified specific populations with genetic susceptibility to bipolar disorder, schizophrenia, ADHD, autism spectrum disorder, and specific traits related to alcohol and tobacco use. Overall, these observations underscore the importance of incorporating cell-type specific manipulations into preclinical research models, in order to better understand the underlying neurobiology of certain neuropsychiatric disorders and develop more targeted treatment options.

In conclusion, this dataset provides comprehensive molecular characterization of cell types in the VTA. The use of snRNA-seq allowed us to investigate the heterogeneity of previously identified cell types, leading to the identification of a potentially novel subset of combinatorial neurons. Furthermore, the ability to assay thousands of genes enabled identification of more selective marker genes, which may be used for more accurate viral targeting for functional studies. Taken together, this dataset will be an essential resource for the greater neuroscience community, but specifically for those studying transcriptionally distinct neuronal subtypes.

## METHODS

### Subjects

Male and female Sprague-Dawley rats (200-250g) were purchased from Charles River Laboratories and individually housed for 1 week prior to tissue collection. Rats were maintained on a 12 h light/dark cycle with ad libitum access to food and water, and were briefly handled for 3 days prior to tissue collection (1-2min/day) to acclimate them to the experimenters. All experimental procedures were approved by the University of Alabama at Birmingham Institutional Animal Care and Use Committee, and were conducted in accordance with the National Institutes of Health Guide for the Care and Use of Laboratory Animals.

### Tissue collection from adult VTA

Rats were euthanized by live decapitation, and brains were rapidly removed and chilled in ice cold Hibernate A media without Ca^2+^ or Mg^2+^ (BrainBits, HACAMG500) for 30-60 sec. Chilled brains were then blocked into 1-mm-thick coronal sections on wet ice. Sections containing the VTA (ranging from -4.92-6.46 anterior-posterior (AP) to Bregma) were transferred to glass petri dishes on dry ice, and the VTA was microdissected from neighboring brain regions (n = 6 rats per sex). Dissected VTA tissue was then transferred to ice-cold centrifuge tubes, and stored at ™80°C until the day of snRNA barcoding and cDNA library preparation.

### Single-nuclei dissociation

Frozen VTA tissue was briefly thawed on wet ice before being chopped by scalpel 100 times in two orthogonal directions. Tissue from three rats per sex was combined and transferred to 5 ml of ice cold lysis buffer (10 mM tris-HCl, 10 mM NaCl, 3 mM MgCl_2_ and 0.1% Igepal in nuclease-free water (Sigma-Aldrich, 18896-50ML) for 15 min, mixing the tissue by inversion every 2 min. The lysis was quenched after 15 min with 5 ml of complete Hibernate A (Thermo Fisher Scientific, A1247501) supplemented with B27, GlutaMAX (Life Technologies, 35050-061), and NxGen RNase Inhibitor (0.2 U/μl; Lucigen, 30281-2). Tissue was then triturated by fire-polished Pasteur pipette (three pipettes of decreasing diameter; 8 to 10 passes per pipette) for 35 min and passed through a 40-μm prewet filter. The samples were then pelleted at 500 rcf for 10 min at 4°C followed by a wash in 10 ml of nuclei wash and resuspension buffer [1× phosphate-buffered saline (PBS), 1% bovine serum albumin (BSA), and NxGen RNase Inhibitor. Supernatant was then removed, and the pellet was gently resuspended in 800 μl of wash buffer before 7-aminoactinomycin D (Thermo Fisher Scientific, 00-6993-50) staining and fluorescence-activated cell sorting (FACS) to further purify the nuclei for sequencing (BD FACS Aria, 70-μm nozzle, BD Biosciences). Immediately after FACS, nuclei were washed a final time at 250 rcf in 10 ml of supplemented Hibernate A containing 1% BSA and RNase Inhibitor for 10 min at 4°C to remove any remaining fine debris and nascent RNA. Nuclei were then brought to a concentration of 1400 nuclei/ μl. A total of 11,367 nuclei pooled from three rats per sex and treatment group were loaded into individual wells of the Chromium NextGem Single Cell Chip (10x Genomics, catalog no. 10000121) using four of the eight available wells.

### snRNA-seq and Analysis

Libraries were constructed according to manufacturer’s instructions for Chromium Next GEM single cell 3’ library and gel bead kit (10x Genomics, v3.1 single index, catalog no. 10000121), which utilizes version 3 chemistry. A total of 21,606 nuclei were captured across 4 GEM (gel bead in emulsion) wells (2 wells per genetic sex) using the 10x Chromium Controller. Each GEM well contained nuclei from 3 separate male or female animals for a total of 6 rats per genetic sex. Libraries were then sequenced on the Illumina NextSeq500 at the Heflin Genomics Core at UAB to an average read depth of ∼26,197 reads per nuclei. A CellRanger reference package was generated using a custom gene transfer file (GTF) from the Ensembl Rn6 genome (version 95) to ensure accurate mapping to both pre-mature, unspliced RNAs as well as mature mRNA. CellRanger filtered outputs were then analyzed with Seurat v3.2.2^85,86^ using R v4.0.2. Nuclei with <200 genes and >5% of reads mapping to the mitochondrial genome were removed from the dataset. Molecular count data from each GEM (gel bead in emulsion) well were then log-normalized with a scaling factor of 10,000. To ensure that the identified clusters were representative of the cell type heterogeneity known to exist in the adult rat VTA, uniform approximation and projection (UMAPs) were generated following data integration for every combination of 21 principal components (10 to 30) and 10 resolution values (0.1 – 1.0). Data from each GEM well were then integrated with FindIntegrationAnchors() and IntegrateData() using 25 principal components and a resolution value of 0.2 as these values produced clusters representative of known cell types in the adult rat VTA. The cell type identity of each cluster was validated by plotting the log-normalized expression values of marker genes for previously identified cell types on top of the final UMAP using FeaturePlot(). The distribution of expression values for these major marker genes were also analyzed with VlnPlot() and DotPlot(). Neuronal subtypes were identified by removing all non-neuronal cells and reclustering with 15 principal components and a resolution value of 0.2. Low confidence clusters with <50 cells were not included in further analyses as they did not represent any specific, transcriptionally defined cell type. Marker genes for the 16 major cell types and 7 neuronal subtypes were identified using a Wilcoxon ranked sum test in which the log-normalized gene expression values for one neuronal subtype were tested against the log-normalized gene expression values of all other neuronal subtypes. Resulting *p*-values were adjusted using the Bonferroni correction based on the number of genes identified in each cluster. Gini coefficients were calculated using the average log23-normalized gene expression values for each cluster with the gini() command from the reldist R package^87,88^. All R code is available at https://gitlab.rc.uab.edu/day-lab.

### MAGMA analysis

Cell type specific gene lists were generated by first identifying marker genes for the 16 transcriptionally distinct cell types expressed in > 10% of cells within the cluster that also had a Bonferroni adjusted p-value < 1×10^−12^ and a log2 fold change value greater than 0.25. Human homologs for these rat genes were then identified by mapping the “HomoloGene.ID” value for each marker gene from rat to human^89^. These “HomoloGene.ID” values can be found at http://www.informatics.jax.org/downloads/reports/HOM_AllOrganism.rpt. Following the generation of these gene sets, Multi-marker Analysis of GenoMic Annotation (MAGMA)^53^ v1.08b was used to identify cell type specific enrichment, and susceptibility, for Alzheimer’s disease^54^, Parkinson’s disease^55^, bipolar disorder^56^, schizophrenia^57^, attention deficit hyperactivity disorder^58^, autism spectrum disorders (ASDs)^59^, and specific traits associated with alcohol and tobacco use^60^. SNPs obtained from summary statistics for each GWAS were mapped to hg19 gene coordinates with a +/-10kb window. Following annotation, these same summary statistics files for each GWAS were then used to perform gene-level analyses with the default snp-wise=mean gene analysis model. Finally, the default competitive gene-set analysis was performed for all gene sets corresponding to the 16 transcriptionally distinct cell types. Resulting empirical p-values were then adjusted with false discovery rate (FDR) for the 16 tests performed for each GWAS.

### Data availability

Sequencing data that support the findings of this study are available in Gene Expression Omnibus (GSE168156). All relevant data that support the findings of this study are available by request from the corresponding author (J.J.D.).

## ACKNOWLEDGEMENTS

We thank all current and former Day Lab members for assistance and support. This work was supported by NIH grants MH114990, DA039650, and DA048348 and the UAB Pittman Scholar Program (JJD), the UAB AMC21 Scholars Program (RAPIII), and a Brain and Behavior Research Foundation Young Investigator Grant (JJT). LI is supported by the Civitan International Research Center at UAB. We thank Shanrun Liu and the UAB Flow Cytometry Core for assistance with 10X Genomics Chromium Capture and FACS, as well as the UAB Heflin Genomics Core facility for assistance with sequencing.

## AUTHOR CONTRIBUTIONS

RAPIII, JJT, SB, and JJD conceived of and performed experiments. RAPIII, JJT, and JJD wrote the manuscript. Data analysis was conducted by RAPIII and JJT, with assistance from LI. JJD supervised all work. All authors have approved of the final version of the manuscript.

---

**Supplementary Figure 1.**
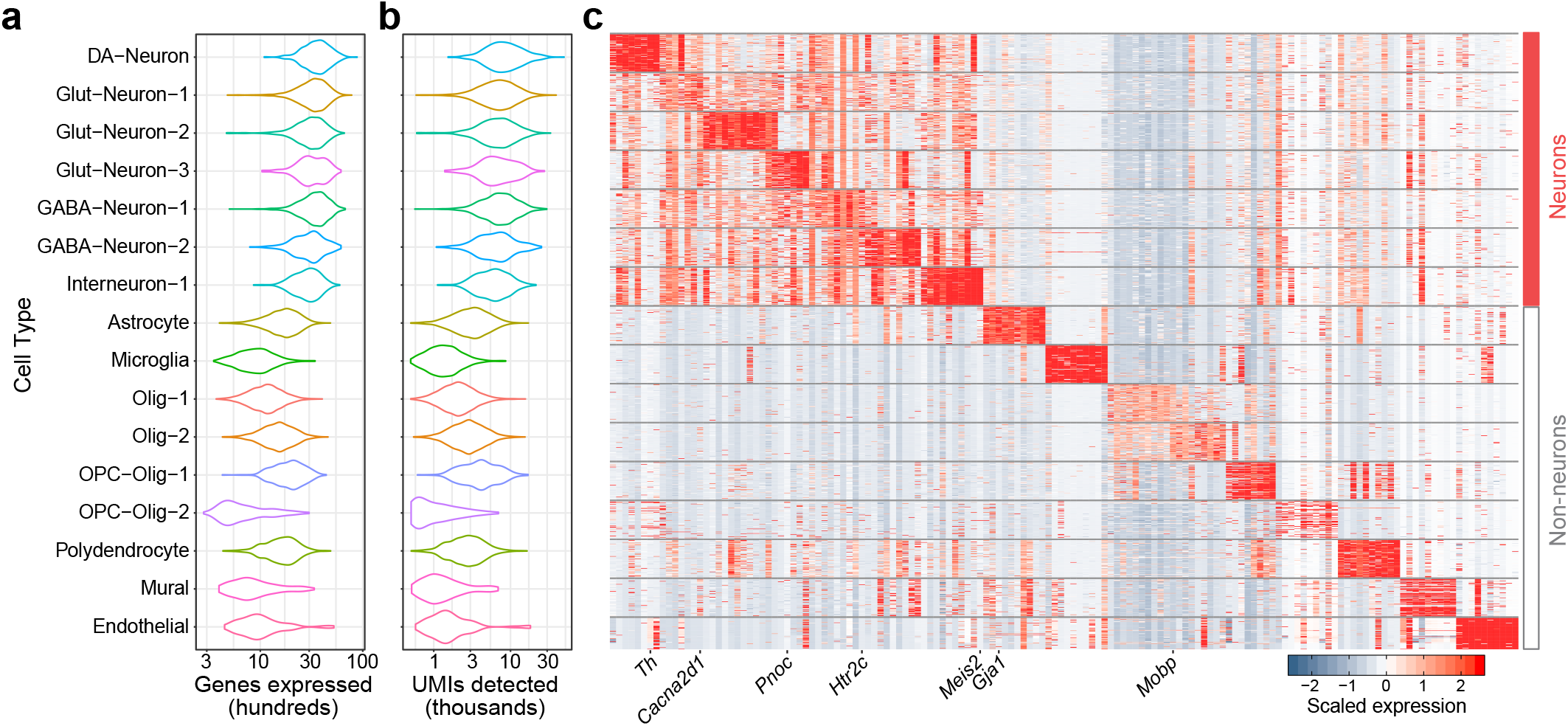
Summary data for snRNA-seq clustering of rat VTA. **a**, Unique genes detected per cell as a function of cell cluster. **b**, Unique molecular identifiers (UMIs) per cell. **c**, Heatmap of top 10 cell-selective marker genes across clusters. Data in heatmap downsampled to 75 cells/cluster.

**Supplementary Figure 2.**
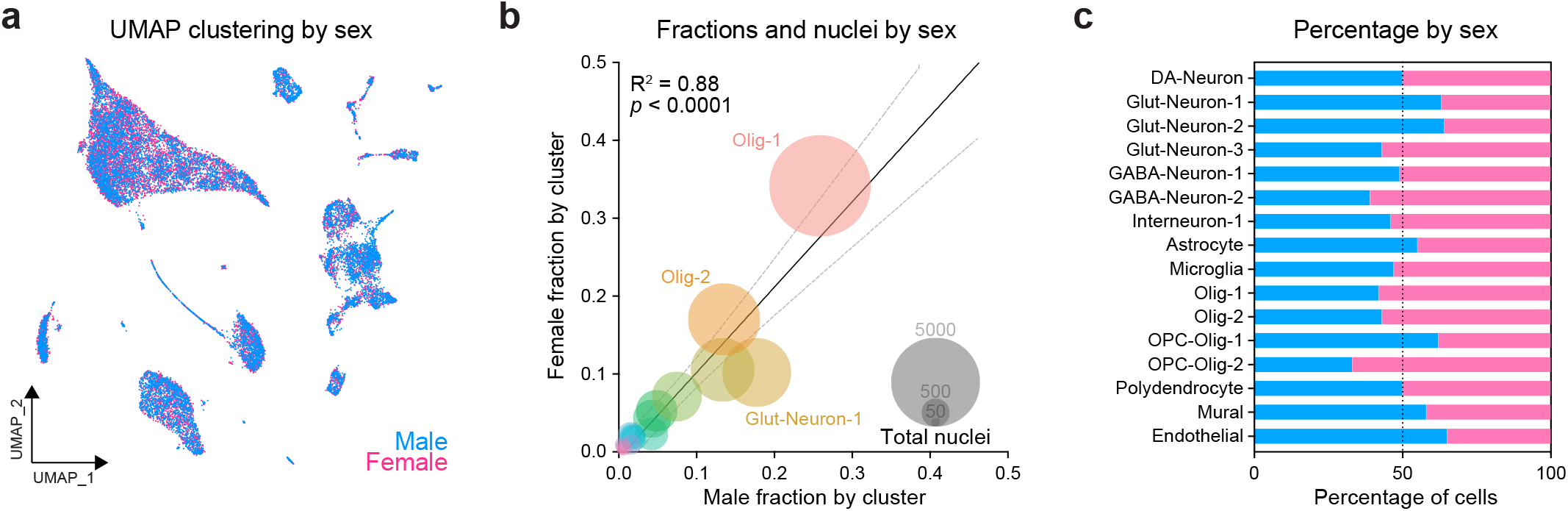
Sex difference comparison of VTA cellular populations. **a**, UMAP showing representation of male and female cells. **b**, Correlation plot of individual cell populations in male and female samples. Size of circle represents the total number of male and female cells in a cluster (combined across sex). Population sizes were highly correlated in male and female samples. **c**, Percentage of cells in each cluster arising from male or female samples reveals similar distibution in each sex.

**Supplementary Figure 3.**
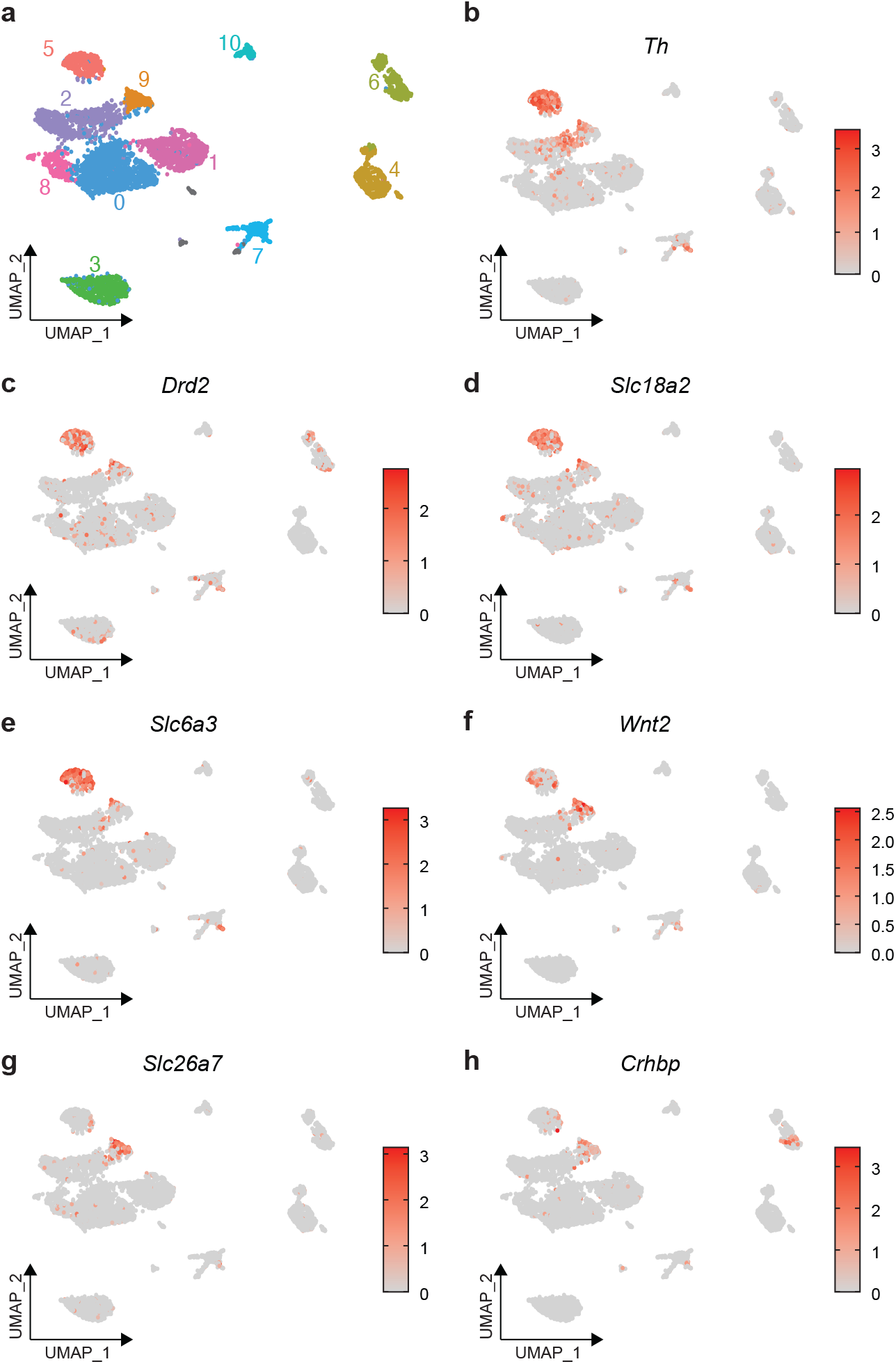
Distribution of expression values for cluster 5 and 9 DEGs within VTA neuronal subclusters. **a**, UMAP depicting location of VTA neuronal subclusters **b-h**. Enrichment of *Th, Drd2, Slc18a2, Slc6a3, Wnt2, Slc26a7*, and Crhbp within VTA neuronal subclusters.

## REFERENCES

1. Volkow, N. D., Fowler, J. S., Wang, G.-J., Swanson, J. M. & Telang, F. Dopamine in drug abuse and addiction: results of imaging studies and treatment implications. Arch. Neurol. 64, 1575–1579 (2007).

2. Savell, K. E. et al. A dopamine-induced gene expression signature regulates neuronal function and cocaine response. Sci. Adv. 6, eaba4221 (2020).

3. Cousins, D. A., Butts, K. & Young, A. H. The role of dopamine in bipolar disorder. Bipolar Disord 11, 787– 806 (2009).

4. Volkow, N. D. et al. Depressed dopamine activity in caudate and preliminary evidence of limbic involvement in adults with attention-deficit/hyperactivity disorder. Arch. Gen. Psychiatry 64, 932–940 (2007).

5. Perez-Costas, E., Melendez-Ferro, M. & Roberts, R. C. Basal ganglia pathology in schizophrenia: dopamine connections and anomalies. J. Neurochem. 113, 287– 302 (2010).

6. Hamilton, P. J. et al. De novo mutation in the dopamine transporter gene associates dopamine dysfunction with autism spectrum disorder. Mol. Psychiatry 18, 1315– 1323 (2013).

7. Day, J. J., Roitman, M. F., Wightman, R. M. & Carelli, R. M. Associative learning mediates dynamic shifts in dopamine signaling in the nucleus accumbens. Nat. Neurosci. 10, 1020–1028 (2007).

8. Saunders, B. T., Richard, J. M., Margolis, E. B. & Janak, P. H. Dopamine neurons create Pavlovian conditioned stimuli with circuit-defined motivational properties. Nat. Neurosci. 21, 1072–1083 (2018).

9. Coddington, L. T. & Dudman, J. T. The timing of action 1. determines reward prediction signals in identified midbrain dopamine neurons. Nat. Neurosci. 21, 1563– 1573 (2018).

10. Kim, H. R. et al. A Unified Framework for Dopamine Signals across Timescales. Cell 183, 1600–1616.e25 (2020).

11. Phillips, P. E. M., Stuber, G. D., Heien, M. L. A. V., Wightman, R. M. & Carelli, R. M. Subsecond dopamine release promotes cocaine seeking. Nature 422, 614–618 (2003).

12. Schultz, W., Dayan, P. & Montague, P. R. A neural substrate of prediction and reward. Science 275, 1593– 1599 (1997).

13. Lerner, T. N., Holloway, A. L. & Seiler, J. L. Dopamine, updated: reward prediction error and beyond. Curr. Opin. Neurobiol. 67, 123–130 (2020).

14. Di Ciano, P., Cardinal, R. N., Cowell, R. A., Little, S. J. & Everitt, B. J. Differential involvement of NMDA, AMPA/kainate, and dopamine receptors in the nucleus accumbens core in the acquisition and performance of pavlovian approach behavior. J. Neurosci. 21, 9471– 9477 (2001).

15. Wise, R. A. Dopamine, learning and motivation. Nat. Rev. Neurosci. 5, 483–494 (2004).

16. Tsai, H.-C. et al. Phasic firing in dopaminergic neurons is sufficient for behavioral conditioning. Science 324, 1080–1084 (2009).

17. Johnson, S. W. & North, R. A. Opioids excite dopamine neurons by hyperpolarization of local interneurons. J. Neurosci. 12, 483–488 (1992).

18. Gysling, K. & Wang, R. Y. Morphine-induced activation of A10 dopamine neurons in the rat. Brain Res. 277, 119–127 (1983).

19. Zell, V. et al. VTA Glutamate Neuron Activity Drives Positive Reinforcement Absent Dopamine Co-release. Neuron 107, 864–873.e4 (2020).

20. Yamaguchi, T., Wang, H.-L., Li, X., Ng, T. H. & Morales, M. Mesocorticolimbic glutamatergic pathway. J. Neurosci. 31, 8476–8490 (2011).

21. Morales, M. & Margolis, E. B. Ventral tegmental area: cellular heterogeneity, connectivity and behaviour. Nat. Rev. Neurosci. 18, 73–85 (2017).

22. Yamaguchi, T., Qi, J., Wang, H.-L., Zhang, S. & Morales, M. Glutamatergic and dopaminergic neurons in the mouse ventral tegmental area. Eur. J. Neurosci. 41, 760– 772 (2015).

23. Yoo, J. H. et al. Ventral tegmental area glutamate neurons co-release GABA and promote positive reinforcement. Nat. Commun. 7, 13697 (2016).

24. Stamatakis, A. M. et al. A unique population of ventral tegmental area neurons inhibits the lateral habenula to promote reward. Neuron 80, 1039–1053 (2013).

25. Granger, A. J., Wallace, M. L. & Sabatini, B. L. Multi-transmitter neurons in the mammalian central nervous system. Curr. Opin. Neurobiol. 45, 85–91 (2017).

26. Kim, J.-I. et al. Aldehyde dehydrogenase 1a1 mediates a GABA synthesis pathway in midbrain dopaminergic neurons. Science 350, 102–106 (2015).

27. Saunders, A. et al. Molecular Diversity and Specializations among the Cells of the Adult Mouse Brain. Cell 174, 1015–1030.e16 (2018).

28. Tiklová, K. et al. Single-cell RNA sequencing reveals midbrain dopamine neuron diversity emerging during mouse brain development. Nat. Commun. 10, 581 (2019).

29. La Manno, G. et al. Molecular diversity of midbrain development in mouse, human, and stem cells. Cell 167, 566–580.e19 (2016).

30. Poulin, J.-F., Gaertner, Z., Moreno-Ramos, O. A. & Awatramani, R. Classification of Midbrain Dopamine Neurons Using Single-Cell Gene Expression Profiling Approaches. Trends Neurosci. 43, 155–169 (2020).

31. Björklund, A. & Dunnett, S. B. Dopamine neuron systems in the brain: an update. Trends Neurosci. 30, 194–202 (2007).

32. Bubar, M. J. & Cunningham, K. A. Distribution of serotonin 5-HT2C receptors in the ventral tegmental area. Neuroscience 146, 286–297 (2007).

33. Schoepp, D. D. Unveiling the functions of presynaptic metabotropic glutamate receptors in the central nervous system. J. Pharmacol. Exp. Ther. 299, 12–20 (2001).

34. Shigemoto, R. et al. Differential presynaptic localization of metabotropic glutamate receptor subtypes in the rat hippocampus. J. Neurosci. 17, 7503–7522 (1997).

35. Muguruza, C., Meana, J. J. & Callado, L. F. Group II metabotropic glutamate receptors as targets for novel antipsychotic drugs. Front. Pharmacol. 7, 130 (2016).

36. Parker, K. E. et al. A Paranigral VTA Nociceptin Circuit that Constrains Motivation for Reward. Cell 178, 653– 671.e19 (2019).

37. Li, X., Qi, J., Yamaguchi, T., Wang, H.-L. & Morales, M. Heterogeneous composition of dopamine neurons of the rat A10 region: molecular evidence for diverse signaling properties. Brain Struct. Funct. 218, 1159– 1176 (2013).

38. Root, D. H. et al. Glutamate neurons are intermixed with midbrain dopamine neurons in nonhuman primates and humans. Sci. Rep. 6, 30615 (2016).

39. Mann, H. B. & Whitney, D. R. On a test of whether one of two random variables is stochastically larger than the other. Ann. Math. Statist. 18, 50–60 (1947).

40. Bauer, M. et al. Delta-like 1 participates in the specification of ventral midbrain progenitor derived dopaminergic neurons. J. Neurochem. 104, 1101–1115 (2008).

41. Christophersen, N. S. et al. Midbrain expression of Delta-like 1 homologue is regulated by GDNF and is associated with dopaminergic differentiation. Exp. Neurol. 204, 791–801 (2007).

42. Pan, H.-X. et al. Gch1 variants contribute to the risk and earlier age-at-onset of Parkinson’s disease: a two-cohort case-control study. Transl Neurodegener 9, 31 (2020).

43. Rudakou, U. et al. Common and rare GCH1 variants are associated with Parkinson’s disease. Neurobiol. Aging 73, 231.e1-231.e6 (2019).

44. Yoshino, H. et al. GCH1 mutations in dopa-responsive dystonia and Parkinson’s disease. J. Neurol. 265, 1860– 1870 (2018).

45. Lewthwaite, A. J. et al. Novel GCH1 variant in Dopa-responsive dystonia and Parkinson’s disease. Parkinsonism Relat. Disord. 21, 394–397 (2015).

46. Gini, C. Measurement of inequality of incomes. The Economic Journal 31, 124 (1921).

47. Rahmati, N., Hoebeek, F. E., Peter, S. & De Zeeuw, C. I. Chloride homeostasis in neurons with special emphasis on the olivocerebellar system: differential roles for transporters and channels. Front. Cell Neurosci. 12, 101 (2018).

48. Molinoff, P. B. & Axelrod, J. Biochemistry of 29. catecholamines. Annu. Rev. Biochem. 40, 465–500 (1971).

49. Daubner, S. C., Le, T. & Wang, S. Tyrosine hydroxylase and regulation of dopamine synthesis. Arch. Biochem. Biophys. 508, 1–12 (2011).

50. Kapatos, G. The neurobiology of tetrahydrobiopterin biosynthesis: a model for regulation of GTP cyclohydrolase I gene transcription within nigrostriatal dopamine neurons. IUBMB Life 65, 323–333 (2013).

51. Kaufman, S. Studies on the mechanism of the enzymatic conversion of phenylalanine to tyrosine. J. Biol. Chem. 234, 2677–2682 (1959).

52. Yang, S. et al. A murine model for human sepiapterin-reductase deficiency. Am. J. Hum. Genet. 78, 575–587 (2006).

53. de Leeuw, C. A., Mooij, J. M., Heskes, T. & Posthuma, D. MAGMA: generalized gene-set analysis of GWAS data. PLoS Comput. Biol. 11, e1004219 (2015).

54. Jansen, I. E. et al. Genome-wide meta-analysis identifies new loci and functional pathways influencing Alzheimer’s disease risk. Nat. Genet. 51, 404–413 (2019).

55. Nalls, M. A. et al. Identification of novel risk loci, causal insights, and heritable risk for Parkinson’s disease: a meta-analysis of genome-wide association studies. Lancet Neurol. 18, 1091–1102 (2019).

56. Stahl, E. A. et al. Genome-wide association study identifies 30 loci associated with bipolar disorder. Nat. Genet. 51, 793–803 (2019).

57. Schizophrenia Working Group of the Psychiatric Genomics Consortium. Biological insights from 108 schizophrenia-associated genetic loci. Nature 511, 421– 427 (2014).

58. Demontis, D. et al. Discovery of the first genome-wide significant risk loci for attention deficit/hyperactivity disorder. Nat. Genet. 51, 63–75 (2019).

59. Grove, J. et al. Identification of common genetic risk variants for autism spectrum disorder. Nat. Genet. 51, 431–444 (2019).

60. Liu, M. et al. Association studies of up to 1.2 million individuals yield new insights into the genetic etiology of tobacco and alcohol use. Nat. Genet. 51, 237–244 (2019).

61. de Jonge, J. C., Vinkers, C. H., Hulshoff Pol, H. E. & Marsman, A. Gabaergic mechanisms in schizophrenia: linking postmortem and in vivo studies. Front. Psychiatry 8, 118 (2017).

62. Brisch, R. et al. The role of dopamine in schizophrenia from a neurobiological and evolutionary perspective: old fashioned, but still in vogue. Front. Psychiatry 5, 47 (2014).

63. Moghaddam, B. & Javitt, D. From revolution to evolution: the glutamate hypothesis of schizophrenia and its implication for treatment. Neuropsychopharmacology 37, 4–15 (2012).

64. Courvoisie, H., Hooper, S. R., Fine, C., Kwock, L. & Castillo, M. Neurometabolic functioning and neuropsychological correlates in children with ADHD-H: preliminary findings. J. Neuropsychiatry Clin. Neurosci. 16, 63–69 (2004).

65. Naaijen, J. et al. Glutamatergic and GABAergic gene sets in attention-deficit/hyperactivity disorder: association to overlapping traits in ADHD and autism. Transl. Psychiatry 7, e999 (2017).

66. Miller, E. M., Pomerleau, F., Huettl, P., Gerhardt, G. A. & Glaser, P. E. A. Aberrant glutamate signaling in the prefrontal cortex and striatum of the spontaneously hypertensive rat model of attention-deficit/hyperactivity disorder. Psychopharmacology 231, 3019–3029 (2014).

67. Madras, B. K., Miller, G. M. & Fischman, A. J. The dopamine transporter and attention-deficit/ hyperactivity disorder. Biol. Psychiatry 57, 1397–1409 (2005).

68. Wu, J., Xiao, H., Sun, H., Zou, L. & Zhu, L.-Q. Role of dopamine receptors in ADHD: a systematic meta-analysis. Mol. Neurobiol. 45, 605–620 (2012).

69. Paul, E. J., Tossell, K. & Ungless, M. A. Transcriptional profiling aligned with in situ expression image analysis reveals mosaically expressed molecular markers for GABA neuron sub-groups in the ventral tegmental area. Eur. J. Neurosci. 50, 3732–3749 (2019).

70. Yamaguchi, T., Sheen, W. & Morales, M. Glutamatergic neurons are present in the rat ventral tegmental area. Eur. J. Neurosci. 25, 106–118 (2007).

71. Kawano, M. et al. Particular subpopulations of midbrain and hypothalamic dopamine neurons express vesicular glutamate transporter 2 in the rat brain. J. Comp. Neurol. 498, 581–592 (2006).

72. Hook, P. W. et al. Single-Cell RNA-Seq of Mouse Dopaminergic Neurons Informs Candidate Gene Selection for Sporadic Parkinson Disease. Am. J. Hum. Genet. 102, 427–446 (2018).

73. Bouarab, C., Thompson, B. & Polter, A. M. VTA GABA neurons at the interface of stress and reward. Front. Neural Circuits 13, 78 (2019).

74. Stuber, G. D., Hnasko, T. S., Britt, J. P., Edwards, R. H. & Bonci, A. Dopaminergic terminals in the nucleus accumbens but not the dorsal striatum corelease glutamate. J. Neurosci. 30, 8229–8233 (2010).

75. Ichinose, H. et al. Hereditary progressive dystonia with marked diurnal fluctuation caused by mutations in the GTP cyclohydrolase I gene. Nat. Genet. 8, 236–242 (1994).

76. Müller, U., Steinberger, D. & Topka, H. Mutations of GCH1 in Dopa-responsive dystonia. J. Neural Transm. 109, 321–328 (2002).

77. Matsui, A. & Williams, J. T. Opioid-sensitive GABA inputs from rostromedial tegmental nucleus synapse onto midbrain dopamine neurons. J. Neurosci. 31, 17729–17735 (2011).

78. De Felipe, C. et al. Altered nociception, analgesia and aggression in mice lacking the receptor for substance P. Nature 392, 394–397 (1998).

79. Zubrzycka, M. & Janecka, A. Substance P: transmitter of nociception (Minireview). Endocr Regul 34, 195–201 (2000).

80. Murtra, P., Sheasby, A. M., Hunt, S. P. & De Felipe, C. Rewarding effects of opiates are absent in mice lacking the receptor for substance P. Nature 405, 180–183 (2000).

81. Ripley, T. L., Gadd, C. A., De Felipe, C., Hunt, S. P. & Stephens, D. N. Lack of self-administration and behavioural sensitisation to morphine, but not cocaine, in mice lacking NK1 receptors. Neuropharmacology 43, 1258–1268 (2002).

82. Korotkova, T. M., Brown, R. E., Sergeeva, O. A., Ponomarenko, A. A. & Haas, H. L. Effects of arousal-and feeding-related neuropeptides on dopaminergic and GABAergic neurons in the ventral tegmental area of the rat. Eur. J. Neurosci. 23, 2677–2685 (2006).

83. Guo, Y., Wang, H.-L., Xiang, X.-H. & Zhao, Y. The role of glutamate and its receptors in mesocorticolimbic dopaminergic regions in opioid addiction. Neurosci. Biobehav. Rev. 33, 864–873 (2009).

84. Schank, J. R. Neurokinin receptors in drug and alcohol addiction. Brain Res. 146729 (2020). doi:10.1016/j.brainres.2020.146729

85. Butler, A., Hoffman, P., Smibert, P., Papalexi, E. & Satija, R. Integrating single-cell transcriptomic data across different conditions, technologies, and species. Nat. Biotechnol. 36, 411–420 (2018).

86. Stuart, T. et al. Comprehensive Integration of Single-Cell Data. Cell 177, 1888–1902.e21 (2019).

87. Relative distribution methods in the social sciences. (Springer-Verlag, 1999). doi:10.1007/b97852

88. Hancock, M. S. Relative Distibution Methods. (CRAN, 2016).

89. Tran, M. N. et al. Single-nucleus transcriptome analysis reveals cell type-specific molecular signatures across reward circuitry in the human brain. BioRxiv (2020). doi:10.1101/2020.10.07.329839

